# Lipidomic and metabolomic profiling on low count human spermatozoa: A robust and reproducible method for untargeted HPLC-ESI-MS/MS-based approach

**DOI:** 10.64898/2026.02.04.703749

**Authors:** Irune Calzado, Manu Araolaza, Mikel Albizuri, Ainize Odriozola, Iraia Muñoa-Hoyos, Iratxe Ajuria-Morentin, Nerea Subirán

## Abstract

1.

**Background:** Human infertility affects approximately 17.5% of the global population, with male factors accounting for nearly half of all cases. The identification of reliable molecular biomarkers is crucial for improving the diagnosis and assessment of male fertility. In this study, we developed and optimized an untargeted high-performance liquid chromatography–electrospray ionization–tandem mass spectrometry (HPLC-ESI-MS/MS) workflow for comprehensive lipidomic and metabolomic profiling of human spermatozoa using only 1.25 million cells per sample.

**Results:** Compared to previous reports, our optimized method achieved unprecedented analytical depth, identifying 473 lipid species and 955 structurally annotated metabolites, corresponding to nearly 7.600-fold improvements in detection efficiency per cell over published approaches. Lipidomic analysis revealed cholesterol, fatty acids, phosphatidylcholines, and phosphatidylethanolamine plasmalogens as the most abundant lipid classes, consistent with the structural complexity of the sperm plasma membrane. Metabolomic profiling showed strong enrichment of lipid-related and steroidogenic pathways, including phospholipid biosynthesis, glycerolipid metabolism and androgen and estrogen metabolism. The integration of lipidomic and metabolomic data highlighted functionally interconnected pathways related to membrane dynamics, energy metabolism, and hormone biosynthesis.

**Conclusions:** Overall, this work establishes a robust, sensitive, and scalable analytical framework enabling high-coverage molecular characterization of spermatozoa from limited sample material, laying the groundwork for future biomarker discovery and clinical applications in male infertility research.

**One Sentence Summary:** Development of a highly sensitive untargeted HPLC-ESI–MS/MS lipidomic and metabolomic workflow that achieves unprecedented molecular coverage from only 1.25 million human spermatozoa, revealing interconnected lipid and metabolic pathways and providing a robust foundation for biomarker discovery in male infertility.

**Graphical abstract:** 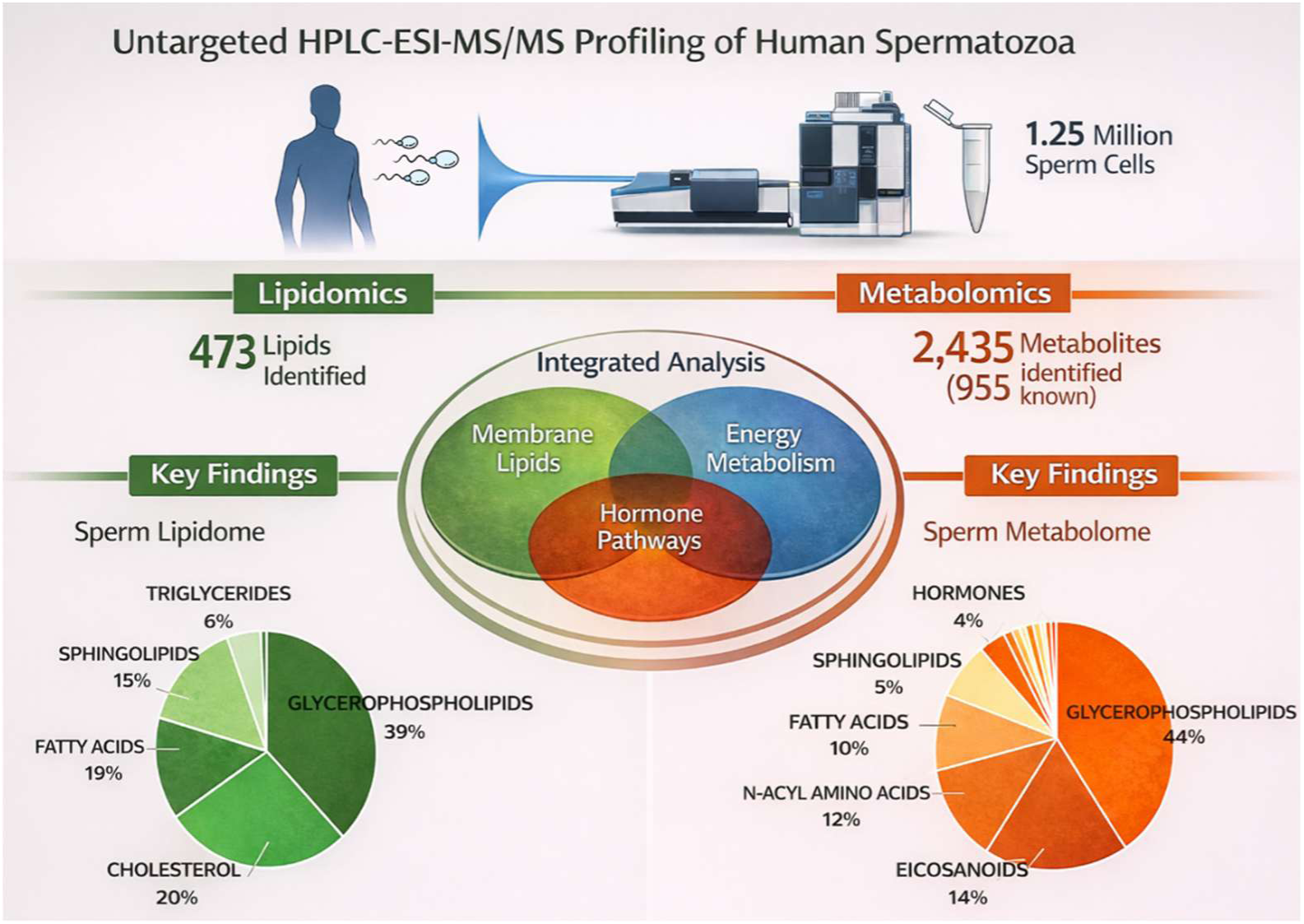

## 2. Background

Human infertility is a global health and social issue, affecting approximately 17.5% of the world population, with male factors contributing to about half of all the cases (1). This highlights a clear and urgent need for reliable biomarkers to better assess male fertility. In this context, advanced omics technologies–including genomics, proteomics, and more recently, lipidomics and metabolomic– have made it possible to find trustworthy indicators of semen quality (2).

In terms of the technique, both nuclear magnetic resonance (NMR) and mass spectrometry (MS)-based techniques have been developed for the study of lipidomics and metabolomics (3). NMR spectroscopy offers several advantages, such as minimal sample preparation and non-destructive analysis of biological samples (4). However, MS-based lipidomics and metabolomics have emerged as the most widely used technique because of its great sensitivity, broader metabolite coverage and capacity to identify small variations in metabolite levels (5,6). Among MS-based platforms, liquid chromatography coupled to mass spectrometry (LC-MS) is currently the most employed technique, although gas chromatography-mass spectrometry (GC-MS), direct infusion MS (DI-MS), and flow injection analysis MS (FIA-MS) have also been applied (7–13). GC-MS is particularly well suited for the analysis of volatile and thermally stable metabolites; however, it often requires chemical derivatization to increase volatility and detectability, limiting its applicability to certain compound classes (3). Therefore, LC-MS has become a popular choice in metabolomics as sample derivatization is not required, and metabolites with more diverse chemical structures and increased molecular sizes can be measured (3,7). On the other hand, DI-MS and FIA-MS offer rapid, high-throughput analysis with minimal chromatographic separation, but are more susceptible to ion suppression and typically provide lower metabolite coverage and structural resolution compared with LC-MS-based approaches (14,15).

To date most of the lipidomic and/or metabolomic studies in the context of male infertility have focused on seminal plasma, a seminal fraction that is essential for sperm viability and for supporting optimal fertilization conditions within the female reproductive tract (16). Accordingly, seminal plasma has been extensively characterized using a variety of analytical platforms, predominantly HPLC-MS/MS (4,17–22), as well as GC-MS (8,9), NMR (23–26) and FIA-MS (10). Conversely, fewer studies have explored the lipidomic and metabolomic profiles of spermatozoa using HLPC-MS/MS (27,28), DI-MS (11), FIA-MS (12), NMR (6) and GC-MS (6,13). This limited number of studies is largely due to the methodological challenges presented by spermatozoa when compared with seminal plasma. For example, in seminal plasma, the protein concentration varies between 35 and 55 g/L, while in spermatozoa is significantly lower, presenting additional challenges for both identification and quantitative analyses (24). The reduced protein abundance in sperm samples often requires the use of a large number of cells to obtain sufficient analytical signal. This is a rather difficult task when cells number is limited, and the quantification needs to be set to a higher threshold level.

Having this in mind, new approaches that allow high-quality extraction from small cell number of spermatozoa are needed. In this study we aimed to develop a reliable and reproducible untargeted HPLC-ESI-MS/MS-based workflow for the comprehensive lipidomic and metabolomic profiling of human spermatozoa. Our results represent an unprecedented increase in lipid and metabolite recovery relative to cell number, establishing a technical foundation for large-scale clinical applications, particularly in scenarios where sperm availability is compromised.

## 3. Materials and Methods

### 3.1 Sample Collection

Semen samples were collected from Galdakao-Usansolo Hospital (Biscay) from male partners of couples undergoing assisted reproductive treatments (IVF/ICSI) at the Assisted Reproduction Units of the Basque Public Health Service (Osakidetza), in accordance with the ethical guidelines approved by the corresponding ethics committee. Eligible participants were between 30 and 50 years of age, had a normal karyotype, no hormonal disorders, no physical abnormalities of the reproductive system, and no history of radiotherapy or chemotherapy. Only samples meeting the inclusion criteria and with signed informed consent were included in the study. Ejaculates were collected following a period of 3 to 5 days of sexual abstinence. Using Computer-Assisted Sperm Analysis (CASA) software, a routine semen analysis was conducted to assess volume, concentration (CON) and progressive motility (PM). Morphology (NF) was assessed by SpermFunc® Diff-Quick Staining Kit. Subsequently, multiple washing steps were conducted to separate seminal plasma from spermatozoa and to eliminate other cell types. Samples were then stored at - 80 °C until further molecular analysis.

Lipidomic and metabolomic extractions were carried out on 10M spermatozoa using a pooled sample with one teratozoospermic, one normozoospermic and one asthenoteratozoospermic sample.

### 3.2 Sample Preparation and Extraction for Lipidomics

Spermatozoa from 3 biological pooled samples (5M cells) were subjected to sonication on a Vibra Cell sonicator (Bioblock Scientific) equipped with a 3 mm A-23 probe with 75% intensity, 7 cycles of 20 seconds ON/20 seconds OFF, and a sperm concentration of 2.5M per aliquot. Each sonicated sample was then divided in order to make 4 technical replicates for lipidomics analysis (**Fig. 1**). Protein concentration for each aliquot was measured using the Micro BCA^TM^ Protein Assay Kit (Thermo Scientific).

**Figure 1.**
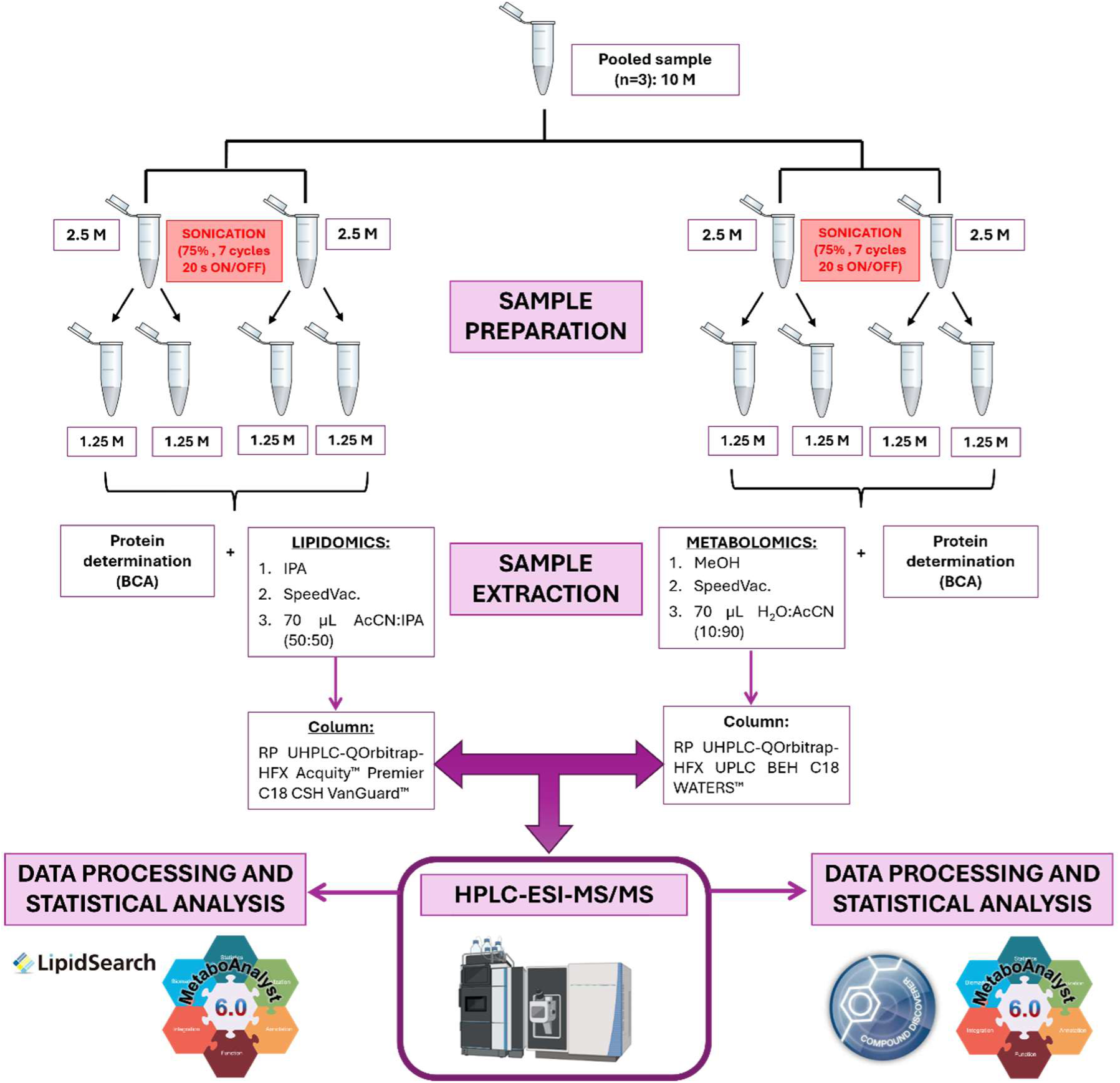
Integrated workflow for lipidomic and metabolomic sample preparation, extraction and analysis of human spermatozoa.

For the lipid extraction, 10 µL of selected internal standard mixtures (Splash™ LipidoMix™, Cer/Sph Mixture I, Cardiolipin Mix I and 24:0(d4) l-carnitine, Avanti Polar Lipids, Inc.) were added, followed by four volumes of isopropanol (IPA) pre-cooled to −20 °C. Samples were then vortexed on a Vortex V-1 plus (biosan) for 1 minute, incubated at room temperature for 10 minutes, and centrifuged at 13.000 × g for 20 minutes at 4 °C. The resulting supernatants were dried using a SpeedVac Concentrator SC250EXP (Thermo Savant) for approximately 2 hours and stored under an inert atmosphere at −80 °C.

### 3.3 Sample Preparation and Extraction for Metabolomics

Spermatozoa from 3 biological pooled samples (5M cells) were subjected to sonication on a Vibra Cell sonicator (Bioblock Scientific) as explained on *Section 3.2* (**Fig. 1**). In parallel to lipidomics, the 4 technical replicates for metabolomics extractions were carried out with 3 volumes of methanol (MeOH) pre-cooled to −20°C. In the same way, samples were then vortexed for 1 minute, incubated at room temperature for 10 minutes, centrifuged at 13.000 × g for 20 minutes at 4 °C, dried and stored under an inert atmosphere at −80 °C.

### 3.4 Lipidomics Analysis by HPLC-ESI-MS/MS

Lipid extracts were analysed using a reverse-phase UHPLC-QOrbitrap-HFX system equipped with an Acquity™ Premier C18 CSH VanGuard™ column (2.1 × 100 mm, 1.7 µm). Columns were pre-equilibrated and cleaned with a water/methanol/formic acid solution (95:5:0.1, v/v) at a flow rate of 100 µL/min. Samples were resuspended in 70 µL of acetonitrile:isopropanol (50:50, v/v), vortexed for 2 minutes and 30 seconds, ultrasonicated for 1 minute, and centrifuged at 13.000 rpm for 4 minutes at 4 °C. Analyses were conducted in both positive and negative electrospray ionization modes (ESI), after injecting 2 µL for ESI+ mode and 5 µL for ESI-mode.

### 3.5 Metabolomics Analysis by HPLC-ESI-MS/MS

Metabolic extracts were analysed using a reverse-phase UHPLC-QOrbitrap-HFX system equipped with an Acquity™ UPLC BEH C18 WATERS™ column (2.1 × 100 mm, 1.7 µm). Columns were pre-equilibrated and cleaned as we explained in the section 3.4. Samples were resuspended in 70 µL of water:acetonitrile (10:90, v/v) and then they were vortexed for 2 minutes and 30 seconds, ultrasonicated for 1 minute, and centrifuged at 13.000 rpm for 4 minutes at 4 °C. Analyses were conducted in both positive and negative electrospray ionization modes (ESI), but in this case the injection volume was 2 µL for both ionization modes.

### 3.6 Data Processing and Statistical Analysis

The raw data obtained for the lipidomic analysis was processed with *Lipid Search^TM^ 4.2.27* software (Thermo Fisher Scientific). The key processing parameters applied were: target database, General; precursor tolerance, 5 ppm; product tolerance, 5 ppm; product ion threshold, 1%; m-score threshold, 2; Quan m/z tolerance, ± 5 ppm; Quan RT (retention time) range, ±0.5 min; use of main isomer filters and ID quality filters A, B, C and D; Adduct ions H^+^, Na^+^ and NH4^+^ for positive ion mode, and H^−^ and HCOO^−^ for negative ion mode. The lipid classes determined were: Acylcarnitine (AcCa), ceramide (Cer), cholesterol (Chol), diacylglycerol (DG), fatty acids (FA), monohexosylceramide (Hex1Cer), dihexosylceramide (Hex2Cer), trihexosylceramide (Hex3Cer), lysophosphatidylcholine (LPC), lysophosphatidylcholine(ether/plasmalogen) (LPCe), lysophosphatidylethanolamine (LPE), lysophosphatidylglycerol (LPG), lysophosphatidylserine (LPS), phosphatidylcholine (PC), phosphatidylcholine(ether/plasmalogen) (PCe), phosphatidylethanolamine (PE), phosphatidylethanolamine(ether/plasmalogen) (PE(e/p)), phosphatidylglycerol (PG), phosphatidylinositol (PI), phosphatidylserine (PS), sphingomyelin (SM), sphingosine (SPH), sulfatide (ST) and triacylglycerol (TG).

After processing, the lipidomics data matrix was cleaned by evaluating correlations between carbon number and the number of double bonds within each lipid class. Linearity was assessed by calculating the coefficient of determination (R²) across different injection volumes, with acceptable values defined at R² > 0.800. Quantification was conducted by normalization of the intensity of the monoisotopic peak of each native lipid species with respect to the intensity of the corresponding monoisotopic peak of the internal standard from Splash LipidoMix, Cer/Sph mixture I, Cardiolipin Mix I and 24:0(d4) l-carnitine (Avanti Polar Lipids, Inc.). Final values were then normalized to protein concentration, expressed in nmol/mg of protein. Lastly, Enrichment Analysis was carried out in the software *MetaboAnalyst 6.0* using the SMPDB database (The Small Molecule Pathway Database).

For metabolomics, Compound Discoverer 3.2 (Thermo Fisher Scientific) was used to process the raw data following the *untargeted metabolomics with statistics detect unknowns and ID using online databases* workflow (*Predicted Compositions, mzCloud Search, Metabolica Search, ChemSpider Search* y *MassList Search*). After processing, the initial data matrix was filtered based on the database match results, applying the following criteria: “*Full Match*,” “*No Results*,” “*Not the Top Hit*,” and “*Partial Match*.” From the resulting data matrix, further cleaning was performed by calculating the coefficient of determination (R²) across different injection volumes, with acceptable values defined at R² > 0.800. Finaly, the (m/z) values were normalized to mg of protein (expressed in (m/z)/mg protein). In the same way, Enrichment Analysis was carried out in the software *MetaboAnalyst 6.0* using the SMPDB database (The Small Molecule Pathway Database).

## 4. Results

### 4.1 Lipidomic Profiling of Human Spermatozoa

Untargeted lipidomic analysis revealed a broad range of lipid species present in human spermatozoa, identified in both positive (ESI+) (**Supp. Table 1A**) and negative (ESI-) (**Supp. Table 1B**) ionization modes. Precisely, we identified 459 lipids in ESI+ and 135 in ESI- and after merging the two datasets, a unified lipid profile of 473 compounds was obtained (**Fig. 2A; Supp. Table 1C**).

**Figure 2.**
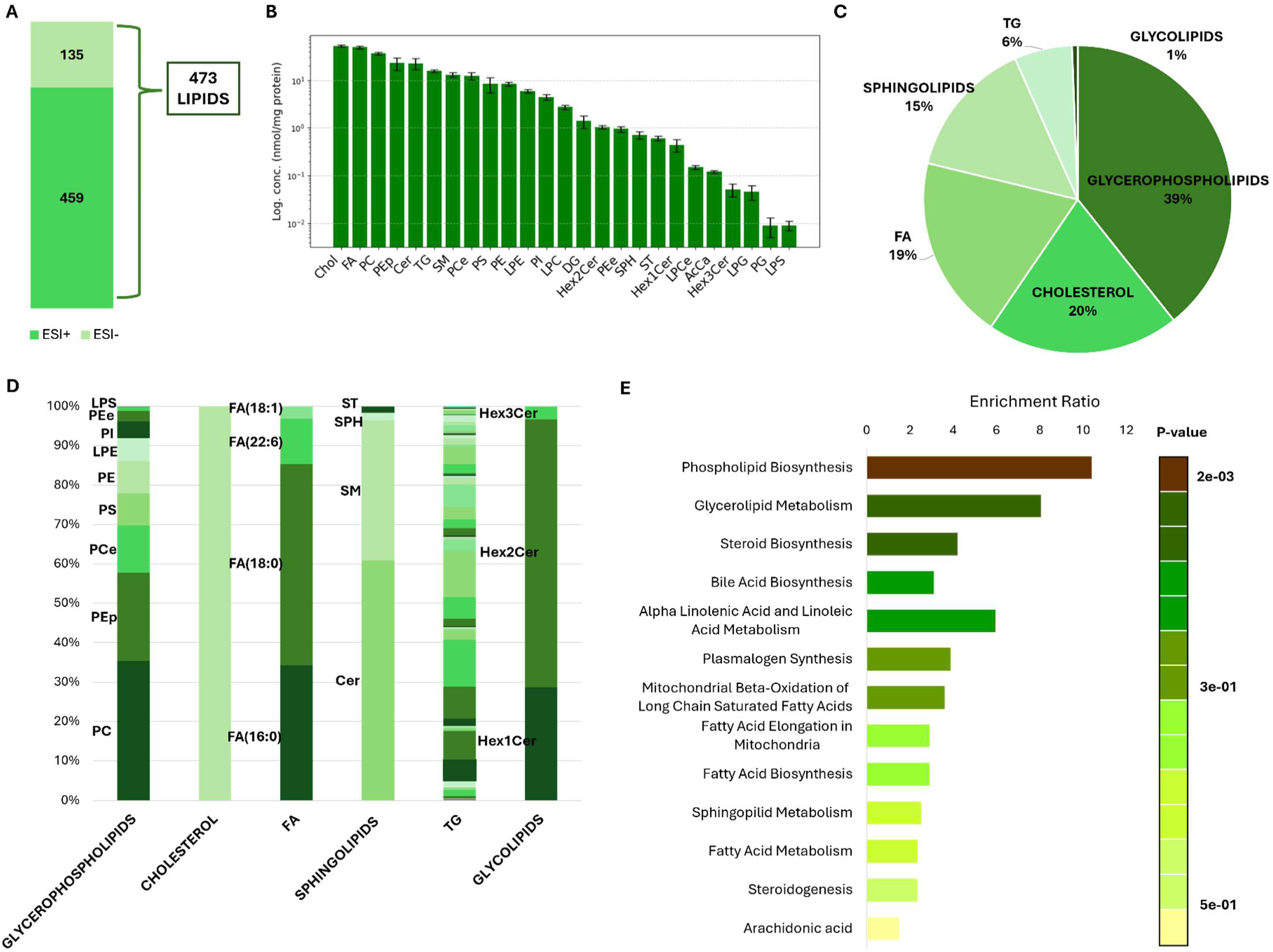
Lipidomics overview in human spermatozoa. (**A)** Feature detection in spermatozoa by untargeted lipidomics. The bar plots show the number of lipids identified in positive (ESI+) and negative (ESI-) ionization modes. The datasets were merged, and the unified dataset with 473 lipids was retained for further analyses. (**B)** Lipid class concentrations in spermatozoa, normalized to protein concentration (nmol/mg protein) and expressed in logarithmic scale (Acylcarnitine (AcCa), ceramide (Cer), cholesterol (Chol), diacylglycerol (DG), fatty acids (FA), monohexosylceramide (Hex1Cer), dihexosylceramide (Hex2Cer), trihexosylceramide (Hex3Cer), lysophosphatidylcholine (LPC), lysophosphatidylcholine(ether/plasmalogen) (LPCe), lysophosphatidylethanolamine (LPE), lysophosphatidylglycerol (LPG), lysophosphatidylserine (LPS), phosphatidylcholine (PC), phosphatidylcholine(ether) (PCe), phosphatidylethanolamine (PE), phosphatidylethanolamine(ether/plasmalogen) (PE(e/p)), phosphatidylglycerol (PG), phosphatidylinositol (PI), phosphatidylserine (PS), sphingomyelin (SM), sphingosine (SPH), sulfatide (ST) and triacylglycerol (TG)). (**C-D)** Distribution of major lipid categories identified in spermatozoa, grouped into: glycerophospholipids (LPC, LPCe, LPE, LPG, LPS, PC, PCe, PE, PEe, Pep, PG, PI, PS; 39%), sterols (cholesterol; 20%), fatty acids (FA; 19%), sphingolipids (Cer, SM, SPH, ST; 15%), triacylglycerols (TG; 6%) and glycolipids (Hex1cer, Hex2Cer, Hex3cer; 1%). **(E)** Lipid set enrichment analysis of the lipidomic dataset performed in *MetaboAnalyst* using the Small Molecule Pathway Database (SMPDB). The bar plot displays the top enriched metabolite sets ranked by enrichment ratio

**Table 1.** Summary of previously reported lipidomic studies performed in spermatozoa, including the technique, number of identified lipids, sperm counts, lipids identified per million sperm cells and references (\**The total number of 479 lipids reported includes both techniques; the original publication does not specify the contribution of each individual method*).

| Technique | Identifications | Sperm number (M) | Lipids per million sperm cells | Reference |
| --- | --- | --- | --- | --- |
| HPLC-ESI-MS/MS (HILIC) | 473 | 1.25 M | 378.4 | Our work |
| (NP)-HPLC-MS + RP- HPLC/MRM-MS | 479*<br>Using both techniques | 10 M | 47.9 | Chen *et al.*, 2021 |
| DI-ESI-MS | 399 | 5.46 M (media) | 73.1 | Furse *et al.*, 2022 |
| HPLC-ESI-MS/MS (HILIC) | 209 | 20 M | 10.5 | Guerra-Carvalho *et al.*, 2024 |
| FIA-MS/MS (targeted) | 140/140 | 464 M (media) | 0.3 | Engel *et al.*, 2019 |

To evaluate the efficiency of our workflow, we compared our dataset with previous analytical platforms used for lipidomics profiling of human spermatozoa. Only four studies reporting broad lipid coverage in sperm cells are published to date (**Table 1**), showing high variability in both the number of identified lipids (ranging from 140 to 479 species), and sperm cell counts used for extraction (from 5.46 to over 464 million). Using a combination of normal phase (NP)-HPLC-MS and reverse-phase (RP)-HPLC-MRM-MS, a total of 479 lipids were identified from 10 million sperm cells, although the authors do not specify how many were detected with each technique. Using direct-infusion (DI)-ESI-MS for an average of 5.46 million sperm cells, the authors reported 399 species; and using a targeted FIA-MS/MS approach, the authors reported 140 lipid species from an average of 464 million spermatozoa. Our HPLC-ESI-MS/MS analysis identified 473 lipid species using only 1.25 million sperm cells, indicating the highest lipid coverage reported to date with the lowest cell number. In fact, when we normalized to cell number, our workflow yielded an efficiency of 378.4 lipids per million sperm cells, markedly surpassing the efficiencies reported in previous studies. The combined NP- and RP-HPLC-MS strategy achieved 47.9 lipids per million cells, whereas DI-ESI-MS reached 73.1 lipids per million and HILIC-HPLC-ESI-MS/MS 10.5 lipids per million, indicating that our method is approximately 8-fold, 5-fold, and 36-fold more efficient, respectively. The targeted FIA-MS/MS approach showed the lowest efficiency, with only 0.3 lipids per million spermatozoa, underscoring that our untargeted HPLC-ESI-MS/MS analysis provides more than three orders of magnitude greater lipid identification density per cell.

Next, to assess the robustness of the method and its applicability to samples with limited sperm availability, we evaluated lipid coverage across decreasing sperm concentrations using the same pooled sample. In this context, lipidomic analysis were performed on two serial dilutions of cell concentration, 0.3125 M and 0.625 M, each analysed in four replicates. Across the concentration range tested, the total number of confidently annotated lipids decreased with lower sperm concentrations, with 415, 443 and 473 species detected at 0.3125 M, 0.625 M and 1.25 M, respectively (**Supp. Fig. 1A**). Violin plots of individual lipid concentrations showed comparable distributions with 0.625 M and 1.25 M sperm cells, and a broader asymmetric distribution at 0.3125 M (**Supp. Fig. 1B**). Venn diagrams of the four replicates per condition showed a large central overlap, with most lipid species consistently detected in all replicates at each sperm concentration (**Supp. Fig. 1C**). The lipid overlap of the four replicates was 82.2%, 84.4% and 85.8% for 0.3125 M, 0.625 M and 1.25 M sperm cells, respectively. Although all three sperm concentrations yielded a higher number of lipid identifications than previously reported studies, the 1.25 M concentration provided higher lipid coverage and reproducibility. While reproducibility across replicates did not reach 100%, more likely due to analytical detection limits, this concentration was considered optimal for subsequent analyses. The observed overlap of 85.8% indicates that the majority of the sperm lipidome is consistently captured and suggests that this workflow can be reliably extended to samples with limited sperm availability.

After confirming the robustness and efficiency of the analytical workflow, we next characterized the qualitative composition of the human sperm lipidome. In our dataset, the detected lipids belonged to 25 major classes, represented in **Fig. 2B**, expressed in logarithmic scale and normalized to protein concentration (nmol/mg protein). Overall, cholesterol (Chol) was the most abundant lipid in spermatozoa, followed by fatty acids (FA), phosphatidylcholines (PC), phosphatidylethanolamine plasmalogens (PEp) and ceramides (Cer). Intermediate concentrations were observed for sphingomyelins (SM), triacylglycerols (TG), ether-linked phosphatidylcholines (PCe), phosphatidylethanolamines (PE) and phosphatidylserines (PS). In contrast, lysolipids (LPG and LPS), glycolipids (Hex1Cer and Hex3Cer), acylcarnitines (AcAc) and phosphatidylglycerol (PG) were detected at lower concentrations.

These lipid classes altogether constitute the main classes normally described in plasma membrane lipids (29): glycerophospholipids (LPC, LPCe, LPE, LPG, LPS, PC, PCe, PE, PEe, Pep, PG, PI, PS; 39%), sterols (cholesterol, 20%), sphingolipids (Cer, SM, SPH, ST, 15%), triacylglycerols (TG; 6%) and glycolipids (Hex1cer, Hex2Cer, Hex3cer, 1%) (**Fig. 2C**).

Following this classification, **Fig. 2D** shows the relative distribution within each lipid class. In the glycerophospholipid fraction, PC and PEp represent the most abundant subclasses, followed by PE, PS, PI, LPE, PCe, and LPS. The fatty acid (FA) pool is dominated by saturated palmitic acid (FA(16:0)) and stearic acid (FA(18:0)), with additional contributions from monounsaturated oleic acid (FA(18:1)) and highly unsaturated docosahexaenoic (DHA) acid (FA(22:6)). Among sphingolipids, SM accounts for the largest proportion, with smaller fractions of Cer, SPH, and ST. The TG group shows a similar distribution of the compounds within this group. Glycolipids are mainly represented by Hex1Cer) and Hex2Cer, with a smaller fraction of Hex3Cer.

Enrichment analysis performed on *MetaboAnalyst* using the Small Molecule Pathway Database (SMPDB), showed phospholipid biosynthesis, glycerolipid metabolism, and steroid biosynthesis as the most enriched pathways (**Fig. 2E**). Additional over-represented pathways are bile acid biosynthesis, alpha-linolenic acid and linoleic acid metabolism, plasmalogen synthesis, mitochondrial beta-oxidation of long-chain saturated fatty acids, and fatty acid elongation and biosynthesis. Lower enrichment ratios are observed for sphingolipid metabolism, fatty acid metabolism, steroidogenesis, and arachidonic acid metabolism.

### 4.2 Metabolomic Profiling of Human Spermatozoa

For the untargeted metabolomic approach a total of 715 features were detected in positive mode (ESI+), of which 364 were successfully annotated based on accurate MS/MS spectra and comparison with database sets (*Predicted Compositions, mzCloud Search, Metabolica Search, ChemSpider Search* y *MassList Search*) (**Supp. Table 2A**), while 351 remained unidentified as no matches were found. Similarly, the negative mode (ESI-) yielded 1.765 features, with 636 compounds annotated (**Supp. Table 2B**) and 1.129 remaining as unknowns. To ensure data robustness and biological interpretability, both ionization datasets were merged and curated, retaining only those metabolites confidently identified (with assigned names and valid database matches). A total of 955 compounds were successfully identified (**Fig. 3A; Supp. Table 2C**), and this unified dataset was then used for all subsequent analyses.

**Figure 3.**
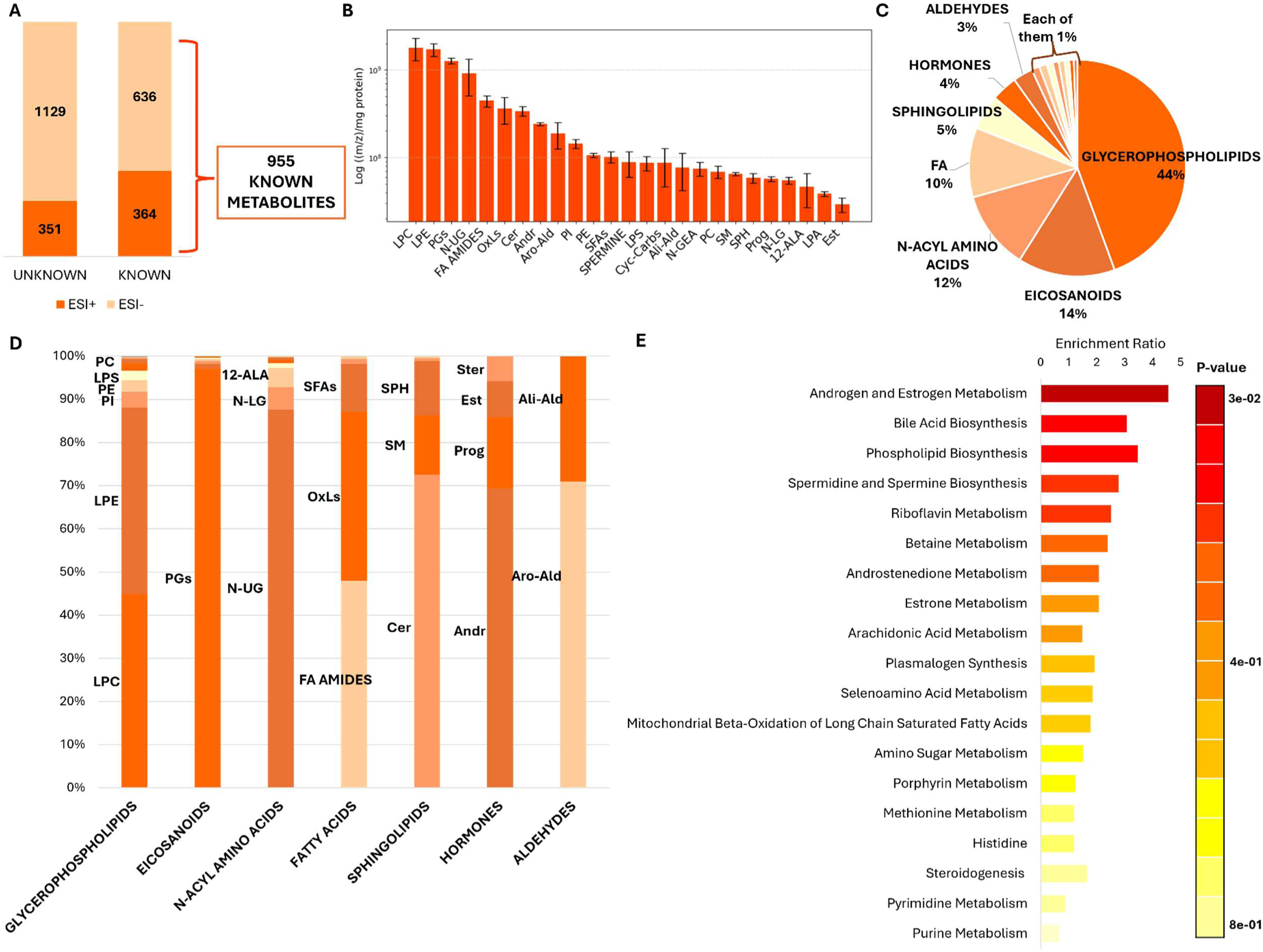
Metabolomics overview in human spermatozoa. (**A)** Feature detection in spermatozoa by untargeted metabolomics. The bar plots show the number of metabolites identified in positive (ESI+) and negative (ESI-) ionization modes, classified as *known* and *unknown*. The datasets were merged, and only the 955 known metabolites were retained for downstream analyses. (**B)** Top 25 metabolites and classes detected in spermatozoa. Relative concentrations are normalized to protein concentration ((m/z)/mg protein) and expressed in logarithmic scale (12-aminolauric acid (12-ALA), androgens (Andr), aliphatic aldehydes (Ali-Ald), aromatic aldehydes (Aro-Ald), ceramides (Cer), cyclic carbohydrates (Cyc-Carbs), estrogens (Est), fatty acid amines (FA amides), lysophosphatidic acid (LPA), lysophosphatidylcholine (LPC), lysophosphatidylethanolamine (LPE), lysophosphatidylserine (LPS), N-gondoylethanolamine (N-GEA), N-lauroylglycine (N-LG), N-Undecanoylglycine (N-UG), oxylipins (OxLs), phosphatidylcholine (PC), phosphatidylethanolamine (PE), phosphatidylinositol (PI), progestogens (Prog), prostaglandins (PGs), saturated fatty acids (SFAs), sphingomyelin (SM), sphingosine (SPH) and spermine). (**C-D)** Classification and distribution of major metabolite categories identified in spermatozoa, grouped mainly into: glycerophospholipids (glycosylphosphatidylinositol (GPI), glycosylphosphatidylglycerol (Glyco-PG), LPA, LPC, LPE, PI, LPS, PC, PE, phosphatidylglycerol (PG), phosphatidylserine (PS); 45%), eicosanoids (docosahexaenoic acid (DHA) derivates, epoxydocosapentaenoic acid derivates (EDE), hydroxyeicosatetraenoic acid derivates (HETE), lypoxines (LX), PGs, thromboxanes (TX); 15%), N-acyl amino acids ((12:0) N-biotinyl, 12-ALA, N-[(4Z,7Z,10Z,13Z,16Z,19Z)-docosahexaenoyl]-L-glutamic acid, N-arachidonyl-L-alanine, N-arachidonoyl-gamma-aminobutyric acid (N-arachidonoylGABA), N-LG, N-Palmitoyl-L-tyrosine, N-UG; 12%), fatty acids (FA amides, monounsaturated fatty acids, OxLs, poly-unsaturated fatty acids (PUFAs), SFAs; 10%), sphingolipids (Cer, glycosphingolipid, lysosphingomyelin (Lyso SM), SM, SPH; 5%), hormonal metabolites (Andr, Est, Prog, steroids (Ster); 4%), aldehydes (Ali-Ald, Aro-Ald; 3%), peptides (1%), amines (1%), carbohydrates (1%), acylcarnitines (1%), amino alcohols (1%), acylcarnitines (1%), bile acids (1%), endocannabinoids (1%) and nucleosides (1%). **(E).** Metabolite-set enrichment analysis performed in *MetaboAnalyst* using the Small Molecule Pathway Database (SMPDB) database. The bar plot represents the top enriched metabolite sets ranked by enrichment ratio.

**Table 2.** Summary of previously reported untargeted metabolomic studies performed in spermatozoa, including the technique, number of identified lipids, sperm counts, metabolites identified per million sperm cells and references.

| Technique | Identifications | Sperm number (M) | Metabolites per million sperm cells | Reference |
| --- | --- | --- | --- | --- |
| HPLC-ESI-MS/MS (HILIC) | 2.435 total<br>955 known | 1.25 M | 764.0 | Our work |
| HPLC-ESI-MS/MS (HILIC) | 112 | 20 M | 5.6 | Guerra-Carvalho *et al.*, 2024 |
| • NMR<br>• GC-MS | 42<br>27 | 150 M<br>15 M | 0.3<br>1.8 | Paiva *et al.*, 2015 |
| GC-MS | 33 | 102,8 M (media) | 0.3 | Zhao *et al.*, 2018 |
| FIA-MS/MS (targeted) | 31/40 | 464 M (media) | 0.1 | Engel *et al.*, 2019 |

Following lipidomic analysis, to also evaluate the efficiency of our metabolomic workflow, we compared our dataset with previous analytical platforms used for metabolomics profiling of human spermatozoa. Published metabolomic studies in human spermatozoa (**Table 2**) have reported far lower numbers of identified metabolites (from 27 to 112 species) despite using markedly higher cell concentration (ranging from 15 million to over 464 million). Specifically, HILIC-HPLC-ESI-MS/MS identified 112 metabolites from 20 million spermatozoa; GC-MS detected between 27 and 33 metabolites from an average of 15 and 102.8 million cells respectively; NMR detected 42 metabolites from 150 million cells, and a targeted FIA-MS/MS workflow reported 31 metabolites using a median of 464 million sperm cells. Our HPLC-ESI-MS/MS analysis identified 2.435 metabolic features, including 955 structurally annotated metabolites, using 1.25 million sperm cells. This also represents the highest metabolomic coverage reported to date. Besides, when considering only structurally annotated compounds, our study achieved an efficiency of 764 metabolites per million sperm cells, indicating a markedly higher metabolite yield per cell than previously reported approaches. In comparison, the HILIC-HPLC-ESI-MS/MS study reported 5.6 metabolites per million spermatozoa, the GC-MS analysis reported 1.8 and 0.3 metabolites per million cells, the NMR analysis reported 0.3 metabolites per million and the targeted FIA-MS/MS method reported only 0.1 metabolites per million, showing that our method provides approximately 140-fold, 430-fold, 2.500-fold, and 7.600-fold higher annotation density per cell, respectively.

We also investigated the impact of spermatozoa concentration on metabolomic coverage by analysing the same pooled sample at reduced cell concentrations (0.3125 M and 0.625 M). In contrast to lipidomics, metabolomic coverage was not affected by sperm concentration, as the total number of confidently annotated metabolites remained constant across all conditions, with 2,435 metabolites detected in each case (**Supp. Fig. 2A**). However, the violin plots of individual lipid concentrations showed similar profiles for 0.625 M and 1.25 M sperm cells, whereas a wider and more asymmetric distribution was evident at the lowest concentration (**Supp. Fig. 2A**). Reproducibility across replicates was high for all conditions. Venn diagram analysis demonstrated complete overlap of metabolite identifications among the four replicates at each sperm concentration (**Supp. Fig. 2A**). Overall, although all three sperm concentrations yielded the same number of metabolites with good reproducibility, variability was lower at 1.25 M sperm cells. Therefore, this concentration was selected for subsequent analyses.

In this context, to characterize the metabolic composition of spermatozoa and given the large number of detected compounds, metabolites were classified into classes. The 25 most abundant classes identified in our dataset are represented in **Fig. 3B**, expressed in logarithmic scale and normalized to protein concentration ((m/z)/mg protein). Overall, lysophospholipids LPC and LPE were the most abundant metabolites in spermatozoa, followed closely by prostaglandins (PGs) and N-undecanoglycine (N-UG), an N-acyl amino acid derivative. Intermediate concentrations were observed for fatty acid (FA) amides, oxylipins (OxLs), ceramides (Cer), androgens (Andr) and aromatic aldehydes (Aro-Ald). In contrast, sphingolipids (SM and SPH), lysophisphatidic acid (LPA), the steroid classes progestogens (Prog) and estrogens (Est), and the N-acyl amino acids N-lauroylglycine (N-LG) and 12-aminolauric acid (12-ALA) were detected at lower concentrations.

The chemical classification of the known metabolites showed that spermatozoa are enriched in lipid-derived molecules, which collectively represented for more than three-quarters of the detected metabolome (**Fig. 3C**). Glycerophospholipids represented the largest group (44%), including multiple subclasses, such as glycosylphosphatidylinositol (GPI), glycosylphosphatidylglycerol (Glyco-PG), LPA, LPC, LPE, PI, LPS, PC, PE, PG, and PS (**Fig. 3D**). Eicosanoids constituted the second major category (14%), comprising DHA derivatives, epoxydocosapentaenoic acids (EDE), hydroxyeicosatetraenoic acids (HETE), lipoxins (LX), thromboxanes (TX), and specially prostaglandins (PGs) which were the most abundant compounds within this family. N-acyl amino acids formed 12% of the metabolome and included several long-chain conjugates such as N-LG, and N-UG. Fatty acids, including saturated, monounsaturated, polyunsaturated, FA amides, and oxylipins, accounted for 10% of the dataset. Additional lipid-related groups included sphingolipids (Cer, GSL, Lyso SM, SM, SPH; 5%) and hormonal metabolites (androgens, estrogens, progestogens, steroids; 4%). Minor metabolite classes were also detected, each contributing 1–3% of the annotated metabolome. These included aldehydes (3%), peptides (1%), amines (1%), carbohydrates (1%), amino alcohols (1%), acylcarnitines (1%), bile acids (1%), endocannabinoids (1%) and nucleosides (1%).

Subsequent enrichment analysis performed on *MetaboAnalyst* showed androgen and estrogen metabolism, bile acid biosynthesis, phospholipid biosynthesis and spermidine and spermine biosynthesis as the most enriched pathways (**Fig. 3E**). Additional enriched pathways also include riboflavin metabolism, betaine metabolism, androstenoedione metabolism, estrone metabolism, arachidonic acid metabolism, plasmalogen synthesis, selenoamino acid metabolism and mitochondrial beta-oxidation of long chain saturated fatty acids. On the other hand, lower enrichment ratios were observed for amino sugar metabolism, porphyrin metabolism, methionine metabolism, histidine, steroidogenesis, pyrimidine metabolism and purine metabolism.

## 5. Discussion

This study presents the optimization of a robust, efficient and reproducible HPLC-ESI-MS/MS-based method for untargeted lipidomic and metabolomic analysis of human spermatozoa. Using an untargeted lipidomic and metabolomic approach with only 1.25 million sperm cells per sample, a total of 473 lipids and 955 metabolites were identified respectively, representing, to our knowledge, the most comprehensive profiling achieved from such a limited number of sperm cells. Although 955 metabolites could be confidently annotated with names in our study, a total of 2.435 features have been detected with their respective retention times and m/z values. This represents, by far, the highest number of metabolite identifications reported in sperm metabolomics to date, surpassing previous studies and highlighting the unprecedented coverage achieved with our workflow.

A critical methodological aspect lies in cell disruption, which can be a hallmark on the increase in our efficiency. We introduced a step of sonication in our procedure prior to analysis, as sonication produces high-frequency ultrasonic waves that generate cavitation events capable of disrupting cell membranes and liberating intracellular compounds without excessive heating or shearing (30). While sonication is becoming increasingly adopted, homogenization remains the procedure of choice in many laboratories (11,27), despite evidence suggesting that sonication produces a greater extraction efficiency in sperm cells than homogenization (31). This approach has been shown to enhance biomolecule extraction in spermatozoa (31), and our data corroborate this, demonstrating that optimized sonication significantly improves both lipid and metabolite coverage. In addition, the high variability on the very few published studies of lipidomics and metabolomics on spermatozoa highlights the current lack of standardization in lipid extraction and quantification methodologies. In most studies conducted on spermatozoa, the cell quantity is not standardized (11–13), and proper normalization is frequently overlooked, making biological interpretation and comparison across studies difficult. Here, we implemented a double-normalization strategy, first by using the same number of cells in all samples and subsequently, by normalizing metabolite abundance to mg of protein. This approach reduced sample-to-sample variability and improved the reliability of quantitative comparisons, ultimately strengthening both lipid and metabolite profiling confidence.

Regarding lipidomics, our results show that membrane lipids are the most abundant lipids in human spermatozoa. Specially, cholesterol represents a key component of the sperm plasma membrane, contributing to its rigidity and stability, while also participating in modulation of the membrane fluidity (32). More precisely, as it was confirmed by the enrichment analysis, cholesterol is the starting point for the biosynthesis of steroid hormones, bile acid biosynthesis and steroidogenesis (33). Glycerophospholipids, on the other hand, are key pathways for membrane assembly and remodelling (34). This is consistent with the enrichment analysis, where it was shown that lipid species from spermatozoa were mainly involved in phospholipid biosynthesis and glycerolipid metabolism. The high enrichment ratios observed for these pathways are consistent with the structural complexity and dynamic nature of the sperm plasma membrane (35).

Fatty acids (FA) were among the most abundant lipid species detected. These molecules are key energy substrates and structural components of complex lipids (36–38) as it could be seen in the enrichment analysis participating in pathways such as plasmalogen synthesis, mitochondrial β-oxidation of long-chain saturated fatty acids, fatty acid elongation, fatty acid biosynthesis and fatty acid metabolism. Besides, spermatozoa showed a predominance of long-chain fatty acids, including polyunsaturated species such as docosahexaenoic acid (DHA, 22:6) and oleic acid (18:1). Long-chain PUFAs incorporated into membrane phospholipids are synthesized from the essential dietary precursors linoleic and α-linolenic acids through elongation and desaturation reactions (39). These long-chain derivatives contribute to maintaining optimal membrane fluidity and flexibility, which are critical during processes such as capacitation, acrosome reaction and sperm motility (35). Moreover, PUFAs serve as precursors for bioactive lipid mediators including prostaglandins and leukotrienes, derivatives of arachidonic acid, also shown in the enrichment analysis (39). Triacylglycerols also showed an importance in the enrichment analysis as they take part in the glycerolipid metabolism and are energy substrates for β-oxidation, through hydrolysis to release fatty acids (27).

Lastly, sphingolipid metabolism, which are involved in membrane signalling and cellular regulation (40,41), is likewise significantly enriched, supporting the functional relevance of the sphingolipids in human spermatozoa.

Metabolomic analysis confirmed the strong representation of lipid-associated biochemical routes. Phospholipid biosynthesis emerged as one of the most significantly enriched metabolite sets, in line with the high proportion of glycerophospholipids detected. On the other hand, the pathway with the highest enrichment ratio was androgen and estrogen metabolism, which can be unified with other detected steroid-related pathways, including steroidogenesis, androstenedione and estrone metabolism. This further supports the presence of metabolites involved in the synthesis and regulation of sex hormones, which are essential for sperm maturation and function (42,43).

Pathways related to fatty acid metabolism (plasmalogen synthesis and mitochondrial β-oxidation of long-chain fatty acids), and their derivatives, such as, arachidonic acid metabolism were also enriched in our results. This reinforces the biological relevance of prostaglandins and other eicosanoids observed in the profiling (39,44), as well as the fatty acids importance as energy substrates.

Notably, Amino acid- and amine- related processes (spermidine/spermine biosynthesis, betaine, selenoamino acid, methionine, histidine and purine metabolism) were also significantly enriched. Interestingly, bile acid biosynthesis was also significantly enriched, despite low absolute abundance. Bile acids are not synthetized directly by spermatozoa but originate mainly from accessory gland secretions such as prostate or seminal vesicles (45,46). Something similar happens with pyrimidine metabolism, which represents nucleosides, determined with a low abundance. Finally, some enriched pathways were not captured by the chemical classification due to their low abundance yet still showed significant pathway enrichment; these include riboflavin metabolism (vitamin B2), amino sugar metabolism, and porphyrin metabolism (represented by a metabolite from the heme group).

Lipidomic and metabolomic profiling provides a detailed view of sperm biochemistry. Lipidomics emphasizes the structural and functional importance of membrane lipids, fatty acids, and sphingolipids, while metabolomics confirms active pathways related to lipid metabolism, steroid biosynthesis, and energy production. The convergence of these datasets demonstrates the robustness of our workflow and links structural lipids to their metabolic context, facilitating interpretation of biological relevance

To sum up, our optimized extraction and MS-based workflow substantially expand current analytical coverage from minimal cellular input, providing a powerful platform for characterizing sperm biochemistry in both research and clinical contexts. Future studies applying this method to large and phenotypically stratified datasets may allow the identification of lipidomic and metabolomic signatures associated with male infertility, ultimately contributing to improved diagnostic accuracy and treatment selection. Beyond providing a high-coverage reference map of the sperm lipidome and metabolome, the method described here offers a scalable analytical framework for future clinical applications. Since only 1.25 million cells are required, this workflow is particularly advantageous for the study of subfertile patients, for whom the availability of sperm is frequently limited. This represents a critical step toward the development of molecular biomarkers capable of diagnosing functional sperm defects, predicting reproductive outcomes, or tailoring personalized therapeutic strategies.

## 6. Conclusions

This study presents an optimized HPLC-ESI-MS/MS workflow that allows comprehensive lipidomic and metabolomic profiling of human spermatozoa from a minimal number of cells. By integrating methodological refinements, including efficient sonication-based extraction and standardized dual normalization, the approach achieved the highest lipid and metabolite coverage reported to date for sperm cells. The resulting datasets reveal that the sperm lipidome is dominated by membrane-related lipids such as cholesterol and glycerophospholipids, which underpin membrane fluidity, integrity, and signalling. Metabolomic analyses further confirmed active biochemical routes associated with lipid metabolism, steroid biosynthesis, and energy production, corroborating the metabolic complexity and functional specialization of spermatozoa. Collectively, these findings demonstrate that untargeted lipidomics and metabolomics can capture molecular processes essential for sperm functionality, offering new opportunities to identify biomarkers predictive of male fertility status. Given its efficiency and scalability, this workflow provides a powerful platform for translational research and future clinical implementation, particularly in the analysis of samples with limited sperm availability, helping bridge the gap between molecular understanding and fertility diagnostics.

## 8 Ethics approval and consent to participate

This study was carried out in accordance with the ethical guidelines approved by the corresponding ethics committee (PI2019184).

## 9 Consent for publication

Not applicable.

## 10 Availability of data and materials

Authors consent to the availability of data and materials. All the data has been uploaded as supplemental information.

## 13 Authors’ contributions

IC, MAL, MAG, AO and IMH carried out the experiments and generated data. IAM provided samples. IC performed the bioinformatic analysis. NS designed the study. NS and IC discussed the data and wrote the manuscript. The authors contributed to the discussion, read and approved the final manuscript.

## 14. Acknowledgements

The authors thank the technical support provided by SGIker of UPV/EHU.

