## Supplementary Data for "Lipidomic and metabolomic profiling on low count human spermatozoa: A robust and reproducible method for untargeted HPLC-ESI-MS/MS-based approach"

Faculty of Medicine and Nursing. University of Basque Country. 48940. Leioa, Bizkaia, Spain.

+34 946015673.

**Keywords:** Male Infertility, Lipidomics, Metabolomics, Human Spermatozoa

**ORCID:**

Irune Calzado: 0000-0002-1787-6792

Manu Araolaza: 0000-0003-1838-4594

Mikel Albizuri: 0009-0008-5484-7414

Ainize Odriozola: 0000-0003-4540-7499

Iraia Muñoa-Hoyos: 0000-0002-5107-6991

Iratxe Ajuria-Morentin:

Nerea Subiran: 0000-0001-9202-1287

Supplementary Figure 1

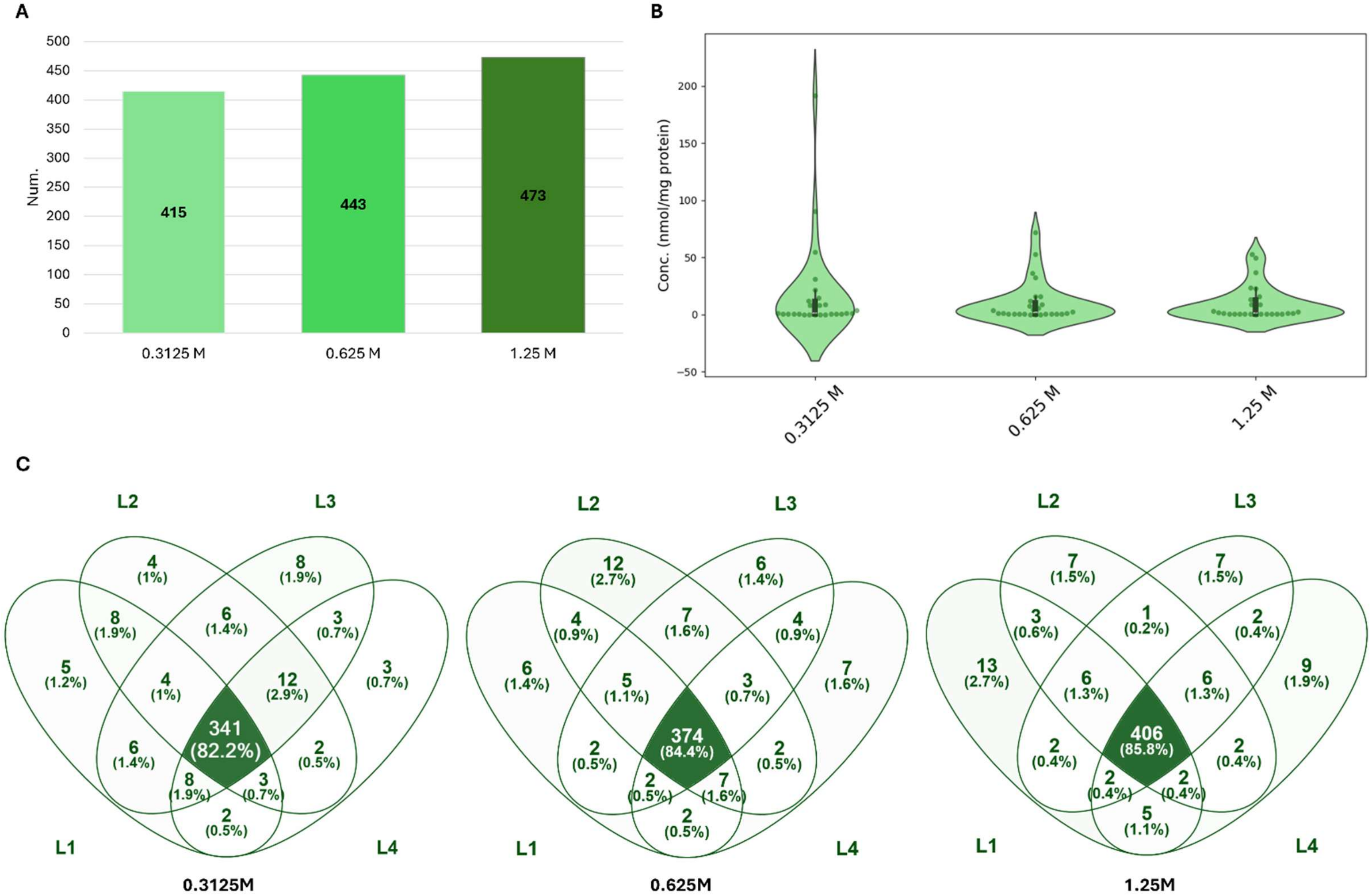

**Supplementary Figure 1. Effect of spermatozoa concentration on lipidomic coverage, variability and reproducibility.** (**A)** Total number of lipids identified in spermatozoa at three different sperm concentrations (0.3125 M, 0.625 M, and 1.25 M). (**B)** Violin plot distribution of lipid concentration at all sperm concentrations. (**C)** Venn diagrams of the overlap of lipid species identified across four replicates (L1-L4) for each spermatozoa concentration. The central intersections represent lipid species consistently detected in all replicates, highlighting analytical reproducibility.

Supplementary Figure 2

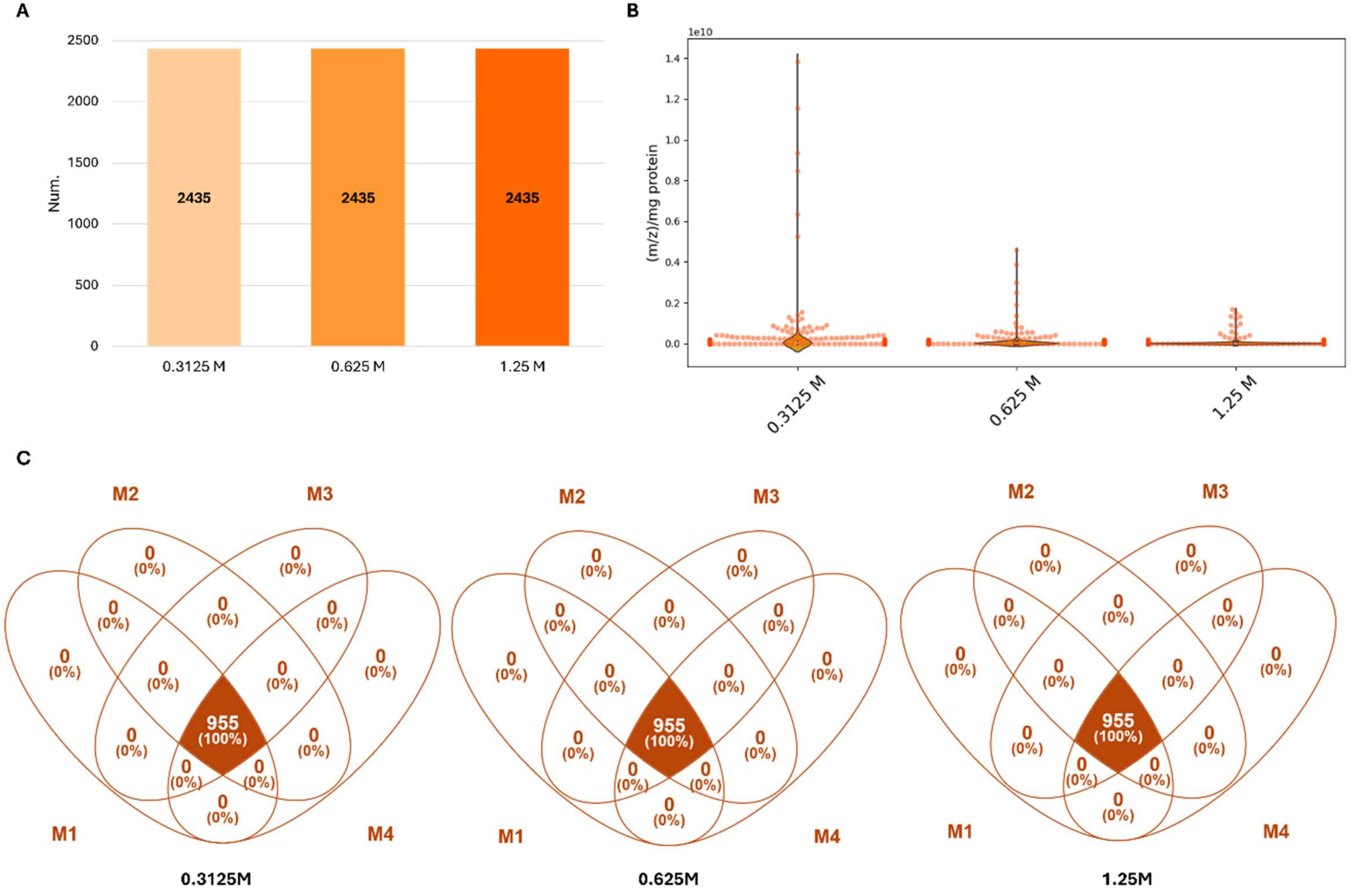

**Supplementary Figure 2. Effect of spermatozoa concentration on metabolomic coverage, variability and reproducibility.** (**A)** Total number of metabolites identified in spermatozoa at three different sperm concentrations (0.3125 M, 0.625 M, and 1.25 M). (**B)** Violin plot distribution of metabolite concentration at all sperm concentrations. (**C)** Venn diagrams of the overlap of metabolite species identified across four replicates (M1-M4) for each spermatozoa concentration. The central intersections represent lipid species consistently detected in all replicates, highlighting analytical reproducibility.

**Supplementary Table S1. Lipidomic analysis dataset. (A)** ESI+ dataset. **(B)** ESI- dataset. **(C)** Unified dataset.

**Supplementary Table S2. Metabolomic analysis dataset. (A)** ESI+ dataset. **(B)** ESI- dataset. **(C)** Unified dataset.
